# A Physiometric Investigation of Inflammatory Composites: Comparison of “A Priori” Aggregates, Empirically-identified Factors, and Individual Proteins

**DOI:** 10.1101/2021.06.21.449259

**Authors:** Daniel P. Moriarity, Lauren M. Ellman, Christopher L. Coe, Thomas M. Olino, Lauren B. Alloy

## Abstract

Most research testing the association between inflammation and health outcomes (e.g., heart disease, diabetes, depression) has focused on individual proteins; however, some studies have used summed composites of inflammatory markers without first investigating dimensionality. Using two different samples (MIDUS-2: N = 1,255 adults, MIDUS-R: N = 863 adults), this study investigates the dimensionality of eight inflammatory proteins (C-reactive protein (CRP), interleukin (IL)-6, IL-8, IL-10, tumor necrosis factor-α (TNF-α), fibrinogen, E-selectin, and intercellular adhesion molecule (ICAM)-1) and compared the resulting factor structure to a) an “a priori”/tau-equivalent factor structure in which all inflammatory proteins equally load onto a single dimension (comparable to the summed composites) and b) proteins modeled individually (i.e., no latent variable) in terms of model fit, replicability, reliability, and their associations with health outcomes. An exploratory factor analysis indicated a two-factor structure (Factor 1: CRP and fibrinogen; Factor 2: IL-8 and IL-10) in MIDUS-2 and was replicated in MIDUS-R. Results did not clearly indicate whether the empirically-identified factor structure or the individual proteins modeled without a latent variable had superior model fit, but both strongly outperformed the “a priori”/tau-equivalent structure (which did not achieve acceptable model fit in any models). Modeling the empirically-identified factors and individual proteins (without a latent factor) as outcomes of medical diagnoses resulted in comparable conclusions. However, modeling individual proteins resulted in findings more robust to correction for multiple comparisons despite more conservative adjustments. Further, reliability for all latent variables was poor. These results indicate that modeling inflammation as a unidimensional construct equally associated with all available proteins does not fit the data well. Instead, individual inflammatory proteins or, potentially (if empirically supported and biologically-plausible) empirically-identified inflammatory factors should be used in accordance with theory.

## Introduction

Atypical inflammatory processes are gaining support as a transdiagnostic feature across many medical and psychiatric conditions. For example, elevated inflammation has been observed in patients with coronary heart disease (Pai et al., 2004), diabetes (Wang et al., 2013), depression (Osimo et al., 2019), and is a proposed mechanism between rheumatoid arthritis (a disease characterized by chronically high inflammation) and secondary adverse medical conditions (e.g., cardiovascular disease; Sattar et al., 2003). Specific immunological theories differ between disease states (e.g., inflammation-induced sickness behavior/depression symptoms to conserve resources and avoid further physical or emotional stress (Dantzer & Kelley, 2007); acceleration of atherogenesis in rheumatoid arthritis leading to heart disease (Sattar et al., 2003)), prompting collection and analysis of inflammatory data in many fields beyond immunology. Given the complexity of this system and its relevance across subfields, it is imperative to critically evaluate current methodologies to ensure optimally-designed and powered studies (Moriarity et al., 2021).

It is common to measure several inflammatory proteins and test them all as independent or dependent variables. Although this facilitates ease of interpretation and can identify protein-specific pathways, maximizing specificity, this approach also results in several complications. First, many theories and hypotheses are described in terms of “inflammation” (e.g., Dooley et al., 2018; Miller et al., 2009; Moriarity et al., 2018; Slavich, 2020), not individual proteins, underscoring a disconnect between theory and analysis. Further, there are concerns about the degree to which any individual protein can be considered a true “biomarker” of inflammation (Konsman, 2019). This approach also invites concerns about the need to adjust for multiple comparisons. To address these concerns, some studies (e.g., Chat et al., 2021; Moriarity et al., 2020; Tait et al., 2019; Vinhaes et al., 2021) have used an “a priori” composite variable consisting of the sum or average of several standardized inflammatory proteins as a measure of general inflammation. There is theoretical rationale for believing that different inflammatory proteins are jointly influenced by a shared, latent, inflammatory process. For example, they are all known to be broadly involved in the initiation and maintenance of, or recovery from, an inflammatory stimulus including tissue damage or infection, sometimes referred to as the inflammatory cascade (Cavaillon & Adib-Conquy, 2002). Additionally, as part of this cascade, many inflammatory proteins are directly involved in the up/down-regulations of others. For example, fibrinogen influences the induction of cytokine/chemokine expression (e.g., IL-6 and TNF-α) via MAC-1 signaling (Amrani, 1990; Davidson, 2013).

However, given the many different component processes of inflammation (e.g., acute phase reaction, upregulation of proinflammatory proteins, activation of the vascular and endocrine systems, neutrophil migration to the site of injury, fibrinolysis, apoptosis, coagulation, and the induction of regulatory anti-inflammatory processes to return to homeostasis (Gruys et al., 2005)), it is plausible that a unidimensional model does not best represent the complexity of this system. Assuming unidimensionality is particularly worrisome with multifaceted processes like inflammation. If different dimensions have different associations with outcomes of interest, aggregating these dimensions could wash out meaningful effects. Additionally, data aggregation uninformed by investigating dimensionality can result in increased measurement error, which, on average, attenuates effect sizes and reduces power (Segerstrom & Boggero, 2020). Further, this approach makes the unlikely assumption that all inflammatory proteins are equally associated with a higher-order inflammatory process (in what is referred to as a “tau-equivalent” model).

It is also important to note that many inflammatory proteins are pleotropic and can contribute to different, even opposing (e.g., pro- and anti-inflammatory), processes under certain conditions. For example, interleukin (IL)-6 typically is described in terms of proinflammatory functions (mediated by classic signaling), but also has some anti-inflammatory functions (mediated by trans-signaling) (Scheller et al., 2011). Further, it is known that the baseline concentrations of various proteins in circulation are partially determined by multiple cellular and tissue sources (i.e., proteins produced by myokines might have higher intercorrelations than proteins produced by different cell types). Consequently, both empirical investigation and careful consideration of biological plausibility are necessary to evaluate the appropriateness and optimal strategies for developing an inflammatory composite (Moriarity, 2021). Indeed, it is important to emphasize that *regardless of empirical support*, inflammatory composite variables should not be used unless they are biologically plausible.

Several published studies have investigated the dimensionality of inflammatory proteins. To our knowledge, only one other study has tested the dimensionality of inflammatory proteins in a non-medical sample. Egnot and colleagues (2018) tested the dimensionality of several inflammatory and coagulatory proteins (C-reactive protein (CRP), IL-6, fibrinogen, intercellular adhesion molecule (ICAM)-1, D-dimer, Lipoprotein (Lp)-a, and pentraxin (PTX)-3). It was concluded that CRP, IL-6, and fibrinogen loaded onto an inflammatory factor and D-dimer and PTX-3 loaded onto a factor indicative of a thrombogenic process and the concomitant vascular perturbation. ICAM-1 and Lp-a did not load onto any observed factors.

A few other studies have conducted factor analyses of inflammatory proteins in medical samples, which are important to compare to the results above to evaluate the generalizability of the dimensionality of inflammation. If the structure of inflammation differs as a function of medical status, this would suggest different inflammatory processes and a need for different methods in different samples. In a sample of acute coronary syndrome patients, Tziakas et al. (2007) tested the dimensionality of CRP, fibrinogen, HDL cholesterol, IL-10, IL-18, and ICAM-1 and found three inflammatory factors: a “systemic inflammation” factor consisting of CRP and fibrinogen, a “local inflammation—endothelial dysfunction” factor consisting of IL-18 and ICAM-1, and an “anti-inflammatory factor” consisting of IL-10 and HDL cholesterol. Koukkunen et al. (2001) conducted an exploratory factor analysis (EFA) on CRP, fibrinogen, IL-6, tumor necrosis factor-α (TNF-α), troponin T, and creatine kinase MB mass in a sample of adults with unstable angina pectoris. This study described two factors: an “inflammation factor” including CRP, fibrinogen, and IL-6, and an “injury” factor including TNF-α, troponin T, and creatine kinase MB mass. Finally, Sakkinen and colleagues (2000) conducted an EFA on 21 different biological characteristics (including several inflammatory proteins) in participants with insulin resistance syndrome. Six biomarkers (fibrinogen, CRP, plasmin-α_2_-antiplasmin, Factor VIIIc, Factor IXc, and fibrin fragment D-dimer) loaded onto what was interpreted as an inflammation factor. HDL cholesterol was tested, but was not found to load onto a factor with other inflammatory markers, as reported by Tziakas et al. (2007).

Some notable similarities between these studies emerge; specifically, 1) CRP, fibrinogen, and IL-6 frequently load onto the same factor (potentially due to their interactive role in the acute phase reaction) when these proteins are included in the dataset and 2) TNF-α and ICAM-1 never loaded onto the same factor as CRP, fibrinogen, and IL-6. It is also worth noting that two of these four studies found multiple inflammation factors, suggesting that inflammation might be best represented as a multidimensional process. Thus, these studies do not support the use of unidimensional composites created by summing standardized values of inflammatory proteins in a dataset (consequently, giving all of the proteins equal weight in the composite) without investigating the empirical structure of the data first. McNeish and Wolf (2020) provide a thorough review of concerns about using sum scores in this manner when the structure of the data is more complex. Additional work must be done to test and replicate the structure of inflammation in populations of interest (e.g., community samples, cancer patients, individuals with depression) to determine the best way to aggregate different inflammatory proteins (and whether this is appropriate). Additionally, none of these studies tested the same panel of inflammatory proteins. Thus, the direct replicability of the structure of an array of inflammatory proteins has never been tested.

### The Present Study

This study sought to critically evaluate a tau-equivalent “a priori” factor structure of inflammatory proteins (comparable to the aggregates used in previously published studies; e.g., Chat et al., 2021; Moriarity et al., 2020; Tait et al., 2019; Vinhaes et al., 2021) by utilizing standard aggregate-creation procedures. First, the structure of eight inflammatory proteins was investigated in a sample of adults. Replicability of this structure was tested in a second sample of similarly-aged adults and model fit was compared to the “a priori”/tau-equivalent model. Structural equation modeling was used in this replication dataset to compare 1) the empirically-identified structure, 2) the “a priori”/tau-equivalent structure, and 3) modeling the inflammatory proteins individually (without a latent variable) as outcomes of several different criterion variables (i.e., heart disease, diabetes, asthma, depression, thyroid disease, peptic ulcer disease, and arthritis) with respect to model fit and results. The authors would like to reiterate that the purpose of this study is not to argue for inflammatory aggregates (doing so requires many different types of data to test characteristics such as structural invariance across acute stress responses, structural invariance across disease states (e.g., illnesses characterized by acute vs. chronic inflammatory abnormalities), and short- and long-term stability). Rather, the goal of this article is to evaluate whether the tau-equivalent “a priori”/tau-equivalent approaches used in the literature can pass several of the preliminary steps of aggregate creation (e.g., replicability, comparability to explorations of factor structure, predictive validity) and to explore alternatives.

## Methods

### Participants and Procedure

Participants were selected from two pre-existing datasets: Midlife in the United States (MIDUS)-2 and MIDUS-Refresher (MIDUS-R). MIDUS-2 (Ryff et al., 2017) consisted of 1,255 (Mage = 55.42 years, 50% female, 78% White) participants between the ages of 25 and 75 who were fluent in English and volunteered to participate in a biomarker collection that included a sera assessment of eight inflammatory proteins (C-reactive protein (CRP), interleukin (IL)-6, IL-8, IL-10, tumor necrosis factor-α (TNF-α), fibrinogen, E-selectin, and intercellular adhesion molecule (ICAM)-1). MIDUS-R (Weinstein et al., 2017) was designed to parallel the procedure of the MIDUS-2 study, and included 863 adults (Mage = 53.53 years, 50% female, 87% white).

### Measures

#### MIDUS-2 and MIDUS-R

##### Inflammatory proteins

Fasting blood draws were collected between 6:00 and 8:30 am for both MIDUS-2 and MIDUS-R. Blood was centrifuged and stored in a −60 °C to −80 °C freezer. Samples were shipped to the MIDUS Biocore Lab on dry ice, where they were stored at −65 °C until assayed. C-reactive protein (CRP) originally was analyzed in plasma via BNII nephelometer (Dade Behring Inc.). Samples falling below the assay range for this method were re-assayed using immunoelectrochemiluminescence using a high-sensitivity array kit (Meso Scale Diagnostics (MSD)). Comparisons of these two methods showed results to be highly correlated. Beginning in 2016, all participants (150 from MIDUS-R) had CRP assayed using the MSD platform. Corrections to account for these changes were applied before the data were made publicly available. Fibrinogen was measured using the same BNII nephelometer. E-Selectin and Intercellular Adhesion Molecule (ICAM)-1 were both measured using ELISA assays (R&D Systems, Minneapolis, MN). Lot-to-lot changes in both E-Selectin and ICAM-1 assays were made throughout the course of the study and adjusted prior to the data being made publicly available to the public. Cytokines (interleukin (IL)-6, IL-8, IL-10, and tumor necrosis factor alpha (TNF-α)) were quantified simultaneously using a MSD V-plex Custom Human Cytokine Kit, and MSD Sector Imager. E-Selectin and ICAM-1 values outside of the detectable range (LLOD = <.1 ng/mL and <45 mg/L, respectively) were set at .09 ng/mL and 44 ng/mL, respectively. MIDUS documentation indicates that none of the other proteins had values outside of the detectable range. Assay ranges and variability for all proteins can be found in the MIDUS documentation available online. Bivariate correlations between the proteins in MIDUS-2 and MIDUS-R are provided in Supplemental Tables 1 and 2, respectively.

###### C-reactive protein

CRP is a pentameric protein, generalized marker of inflammation, and an acute phase reactant upregulated by IL-6 (Davidson, 2013). CRP is primarily synthesized by the liver and can activate the complement system, promoting phagocytosis, and facilitate antibody/antigen binding.

###### Fibrinogen

Fibrinogen is a glycoprotein complex and acute phase protein synthesized in the liver and upregulated by IL-6. It is involved in creating blood clots, regulating thrombin, and influencing leukocyte migration (Amrani, 1990; Davidson, 2013). Additionally, it influences the induction of cytokine/chemokine expression (e.g., IL-6 and TNF-α) via MAC-1 signaling. Breakdown products of fibrinogen (e.g., D-dimers) stimulate the release of several inflammatory proteins including CRP and IL-6 (Davidson, 2013).

###### E-selectin

E-selectin is a selectin cell adhesion molecule expressed by endothelial cells and activated by cytokines. Local release of IL-1 and TNF-α induces over-expression of E-selectin, which then recruits leukocytes to the site of injury (Imhof & Dunon, 1995).

###### Intracellular Adhesion Molecule-1

ICAM-1 increases rapidly in response to TNF-α and IL-1 and influences neutrophil adhesion (Divietro et al., 2001) and recruitment of macrophages. It is expressed by the vascular endothelium, macrophages, and lymphocytes. There is also evidence that ICAM-1 is involved in the secretion of TNF-α (Etienne-Manneville et al., 1999).

###### Interleukin-6

IL-6 is responsible for stimulating acute phase protein synthesis in the liver, production/trafficking of neutrophils and proliferation of B cells (Davidson, 2013; Fielding et al., 2008). It is produced by a wide variety of cells including liver cells, macrophages, osteoblasts, and monocytes (Davidson, 2013). In addition to its pro-inflammatory role, it also has anti-inflammatory properties and is involved in the regulation of TNF-a and IL-10 (Scheller et al., 2011).

###### Interleukin-8

IL-8 (also known as neutrophil chemotactic factor) is a chemokine produced by macrophages and other types of cells (e.g., epithelial cells, smooth muscle cells in the airway, and endothelial cells) (Hedges et al., 2000). It is a key protein promoting neutrophil adhesion (both in terms of adhesion promotion and inhibition) and migration toward injury sites via chemotaxis, and also stimulates phagocytosis (Divietro et al., 2001; Dixit & Simon, 2012; Luscinskas et al., 1992).

###### Interleukin-10

IL-10 is predominantly produced by monocytes and lymphocytes and, primarily, is an anti-inflammatory cytokine that regulates pro-inflammatory proteins (e.g., TNF-a, IL-8, IL-6) as well as enhancing B-cell survival and antibody production (Kessler et al., 2017; Sun et al., 2009).

###### Tumor necrosis factor alpha

TNF-α is a predominantly proinflammatory cytokine primarily released by macrophages (in addition to other cells such as lymphoid cells, adipose tissue, and mast cells) to recruit other cells to activate immune processes (Olszewski et al., 2007). Many proinflammatory functions of TNF-α are apoptotic (promotes programmed cell death) in nature (Gough & Myles, 2020). TNF-α can influence migration of neutrophil to injury sites (Smart & Casal, 1994) and induces ICAM-1 (Burke-Gaffney & Hellewell, 1996).

##### Medical Status

Participants’ medical history was assessed on the day of the study visit via interview. This interview found that 11.6/9.3% of participants with complete biomarker data in these samples (MIDUS-2/MIDUS-R, respectively) reported a history of heart disease, 12.2/10.9% diabetes, 12.4/18.1% asthma, 24.0/34.4% depression, 12.5/12.1% thyroid disease, 43.1/36.4% arthritis, and 5.2/5.2% reported experiencing a peptic ulcer.

### Analyses

All analyses were conducted in R 3.6.2 (R Core Team, 2013). Several proteins were substantially skewed in both datasets. Instead of modifying the data via transformation we elected to use modeling procedures robust to non-normality (described below), in line with best practices (Wilcox & Rousselet, 2018)..

#### Exploratory Factor Analysis

Parallel analysis was used to determine the number of factors to retain using *EFA.dimensions* (Connor, 2020). Unlike other factor retention methods, parallel analysis allows for correction for the effects of sampling error. Eigenvalues were generated using permutations of the raw dataset to create eigenvalues that could be compared to the eigenvalues produced by the factor analysis on the dataset. When an eigenvalue generated from the factor analysis is higher than that generated from the parallel analysis, it can be assumed that the eigenvalue represents a real factor that accounts for more variance than a factor based on the permutated data (Horn, 1965). Thus, the number of factors retained was determined by the number of eigenvalues for which the eigenvalue for the raw data is higher than the eigenvalue that correspond to the 95th percentile of the distribution of the permutated data eigenvalues (consistent with *EFA.dimensions* defaults). Exploratory factor analyses were conducted using *lavaan* in MIDUS-2 using the Geomin rotation to allow for correlated factors (Rosseel, 2012). Models were estimated using maximum likelihood estimation with robust standard errors (MLR) to account for non-normality of the inflammatory proteins. Complex loadings (i.e., items loading onto more than one factor) would be allowed due to the pleotropic nature of these biomarkers. Loadings at or above .30 were considered to load onto a factor. Given that, unlike the creation of self-report items for a survey, it is unlikely and inadvisable that researchers remove proteins that don’t load onto a latent factor from the dataset, there was no iterative process of dropping proteins below this threshold and re-running the parallel analysis and EFA.

#### Confirmatory Factor Analysis/Structural Equation Modeling

Confirmatory factor analyses (CFAs) and structural equation models (SEM) were estimated in MIDUS-R for the factor structure found in the EFA from MIDUS-2. All models were conducted in *lavaan* (Rosseel, 2012). Models were estimated using maximum likelihood estimation with robust standard errors (MLR) to account for non-normality of the inflammatory proteins. The variance of latent variables was set to 1, all latent means set to zero, and all factor loadings, unless otherwise noted in the results, were freely estimated. Next, model fit of the empirically-identified factor structure was compared to the “a priori”/tau-equivalent unidimensional factor structure with the factor loadings of all proteins constrained to equality. In addition to interpretation of standard goodness-of-fit indices (i.e., good fit indicated by comparative fit index [CFI] ≥ .95, root mean square-error of approximation [RMSEA] ≤ .06, and standardized root-mean-square residual [SRMR] ≤ .08, and non-significant chi-square test; Hu et al., 2009), AIC and BIC will be compared between models. Unlike most other fit indices, AIC and BIC can be directly compared, with lower values indicating preferable model fit. Additionally, it is important to note that the chi-square test of significance is over-powered in large sample sizes (Bollen, 1989). SEMs also were conducted in MIDUS-R comparing the empirically-identified structure, “a priori”/tau-equivalent structure, and individual inflammatory proteins as outcomes of several criterion variables (heart disease, diabetes, asthma, tuberculosis, thyroid disease, peptic ulcer disease, and arthritis). Model fit and results were compared.

#### Reliability

Factor reliability was quantified via coefficient ω (Raykov, 2001) using the R package *semTools* (Jorgensen et al., 2021) in MIDUS-2. Coefficient ω was selected over Cronbach’s α because a) ω makes fewer and more realistic assumptions compared to α and b) problems with misestimation of reliability are far less likely. For a more in-depth argument for the adoption of ω over α as the new field standard refer to Dunn et al. (2014).

## Results

### Exploratory Factor Analysis

The exploratory factor analysis was conducted using the 1231 complete observations out of the total 1255 blood draws in MIDUS-2 (some participants had missing data for some proteins and the models estimated can’t handle missing data). The parallel analysis (Table 1) indicated two factors (i.e., two factors had eigenvalues greater than the corresponding factors in the random data). The resulting EFA results can be found in Table 2. CRP and fibrinogen loaded onto Factor 1 and IL-8 and IL-10 loaded onto Factor 2. No other proteins had a loading > .3 on either factor. These factors accounted for 14% and 11% of the total variance, respectively. In comparison, a single factor model only accounted for 15% of the total variance.

**Table 1.** Parallel Analyses.

| Factor # | Eigenvalues<br>Real Data | Eigenvalues<br>95<sup>th</sup> % Random Data |
| --- | --- | --- |
| 1 | 1.905 | 1.169 |
| 2 | 1.302 | 1.101 |
| 3 | 1.050 | 1.059 |
| 4 | .988 | 1.028 |
| 5 | .941 | 1.004 |
| 6 | .715 | .980 |
| 7 | .625 | .956 |
| 8 | .474 | .922 |

Note: N = 1231

**Table 2.** Exploratory Factor Analysis (EFA) in MIDUS-2.

|  | Factor 1 | Factor 2 |
| --- | --- | --- |
| C-reactive Protein | .77* | -.01 |
| Interleukin-6 | .10 | .06 |
| Tumor Necrosis Factor-α | .22 | .28 |
| Interleukin-8 | -.04 | .76* |
| Interleukin-10 | .12 | .43* |
| Fibrinogen | .63* | -.02 |
| Intercellular Adhesion Molecule-1 | .21 | .12 |
| E-selectin | .17 | .12 |
| Correlation Between Factors | .10 |  |
| Proportion of Variance Explained | .14 | .11 |
| Proportion of Variance Explained by Single Factor Solution | .15 |  |

Note: N = 1231, \* = substantially loaded onto the factor

In the empirically-identified structure, the inflammatory factor including CRP and fibrinogen was interpreted to reflect shared variance between these proteins involved in acute phase reaction processes localized in the liver. Both CRP and fibrinogen are acute phase reactants synthesized in the liver and upregulated in response to acute stress and inflammatory challenges (Amrani, 1990; Davidson, 2013). Further, breakdown products of fibrinogen (e.g., D-dimers) can upregulate CRP, creating a positive feedback loop (Davidson, 2013). The second inflammatory factor, consisting of IL-8 and IL-10 was interpreted to reflect shared variance between these proteins related to inflammatory processes (both pro- and anti-inflammatory) particularly associated with neutrophil activity (Korthuis et al., 1994). IL-8 can influence neutrophil adhesion (Divietro et al., 2001), and stimulate neutrophils to migrate to injury sites (Dixit & Simon, 2012). IL-10 serves as an important mediator of neutrophil activity via regulation of proinflammatory cytokines and CXC keratinocyte-derived chemokine molecules (e.g., IL-8; Kessler et al., 2017; Sun et al., 2009). Although commonly described as a “proinflammatory cytokine”, like IL-10, IL-8 also has anti-inflammatory functions (Qazi et al., 2011).

### Confirmatory Factor Analyses

The confirmatory factor analyses were conducted using the 849 complete observations out of the total 853 total blood draws in MIDUS-2 (some participants had missing data for some proteins and the models estimated can’t handle missing data). First, the factor structure described above was re-estimated in MIDUS-2 (Figure 1a). Typically, latent variables need three indicators to be properly identified in these models. Given that each factor only had two indicators, the decision was made to constrain the factor loadings of CRP on Factor 1 and IL-8 on Factor 2 to equality to free up degrees of freedom. These two loadings were chosen because they were almost identical in the original EFA (.77 and .76, respectively). Model fit for replication of the empirically-identified structure had mixed support for its replicability. Both the robust CFI and SRMR met conventional criteria for acceptable fit (robust CFI = .966, SRMR = .022). RMSEA was slightly above the traditional cut-off of .06 (robust RMSEA = .068, 90% CI = .042-.097). The chi-square test was significant (robust χ^2^(2) = 19.101, *p* < .001), but this may be uninformative with sample sizes this large. Additionally, the EFA was well-replicated at the factor loading level. Three of the four originally estimated factor loadings were between the 95% confidence intervals for the factor loadings in the replication model (Table 4).

**Figure 1a.**
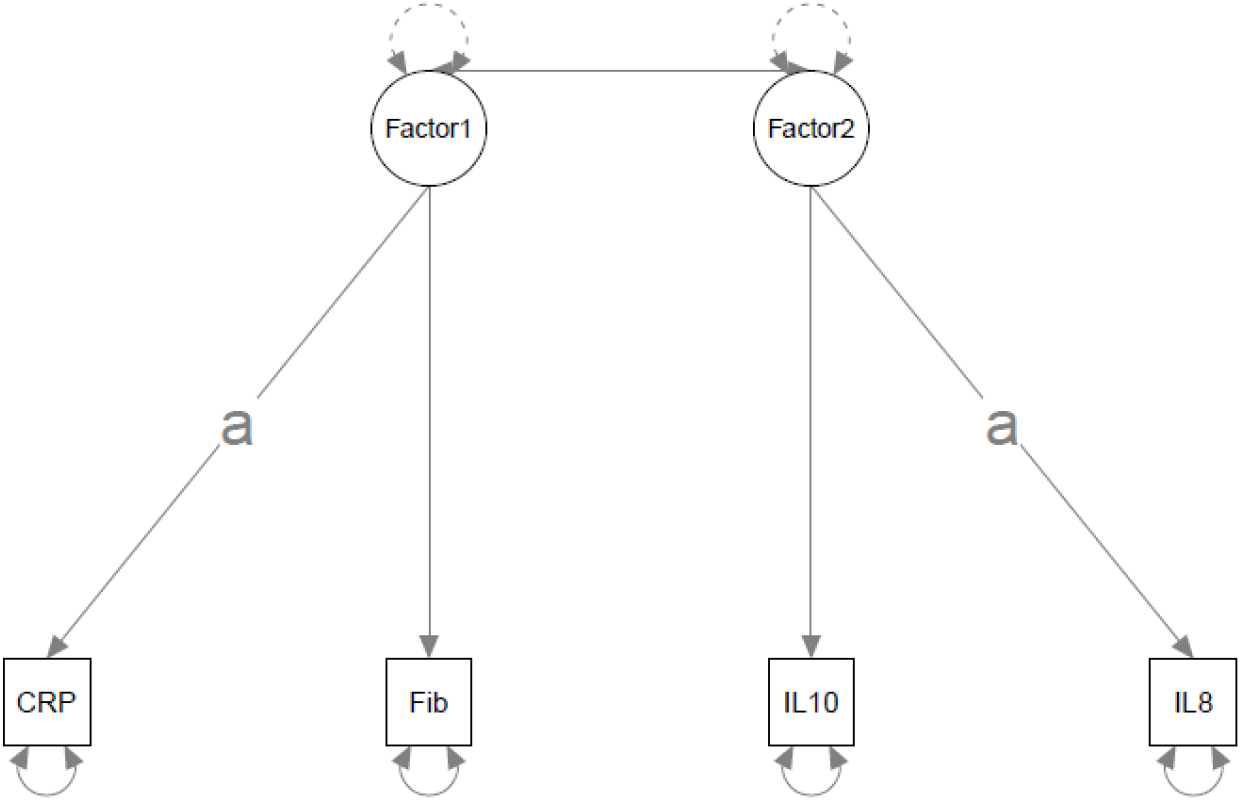
Empirically-identified Structure.

Second, this empirically-identified factor structure was compared to the “a priori”/tau-equivalent factor structure with all proteins having an equal weight on a single factor (Figure 1b). Model fit for the “a priori”/tau-equivalent model was poor: robust CFI = .086, robust RMSEA = .184 (90% CI = .163-.206), and robust SRMR = .160. The chi-square test was significant (χ^2^(27) = 246.150, *p* < .001). Because there are a different number of proteins included in these two models, comparison of AIC/BIC statistics would be inappropriate. According to standard interpretive thresholds, reliability was poor for all inflammatory factors. In order of best to worst: Factor 1 (ω = .351), Factor 2 (ω = .298), and the “a priori”/tau-equivalent model (ω = .001).

**Figure 1b.**
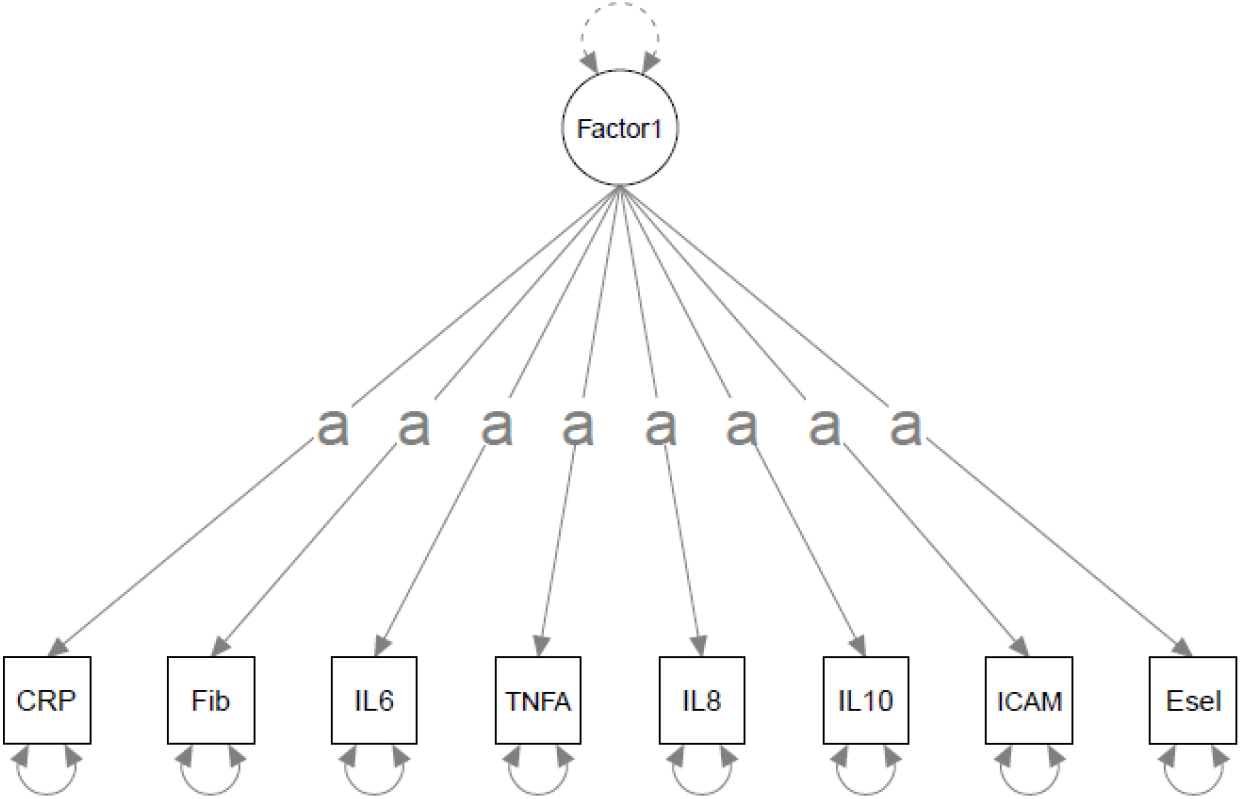
A Priori/Tau-Equivalent Structure. Note: “a” denotes loadings constrained to equality, CRP = C-reactive Protein, Fib=fibrinogen, IL = interleukin, TNFA= Tumor Necrosis Factor-α, ICAM = Intracellular Adhesion Molecule-1, Esel = E-selectin.

### Health Criterion Models/Predictive Validity

To evaluate model fit and associations with external criteria, several health conditions associated with inflammation (heart disease, diabetes, asthma, depression, thyroid disease, peptic ulcer disease, and arthritis) were modeled as predictors of a) the empirically-identified factors and the individual proteins not included in these factors (Figures 2a), b) the “a priori”/tau-equivalent factor (Figure 2b), and c) individual proteins (modeled without a latent variable; Figure 2c) in separate models. Model fit for the empirically-identified model was good (robust CFI = .956, robust RMSEA = .053 (90% CI = .043-063), robust SRMR = .021). As expected with this sample size, the chi-square tests was significant (robust χ^2^(23) = 109.019, *p* < .001). Similar to the results of the CFA, model fit of the “a priori”/tau-equivalent model was unacceptable: robust CFI = .077, robust RMSEA = .133 (90% CI = .126-.139), and robust SRMR = .105. The chi-square test was significant (χ^2^(76) = 1231.540, p < .001).

**Figure 2a.**
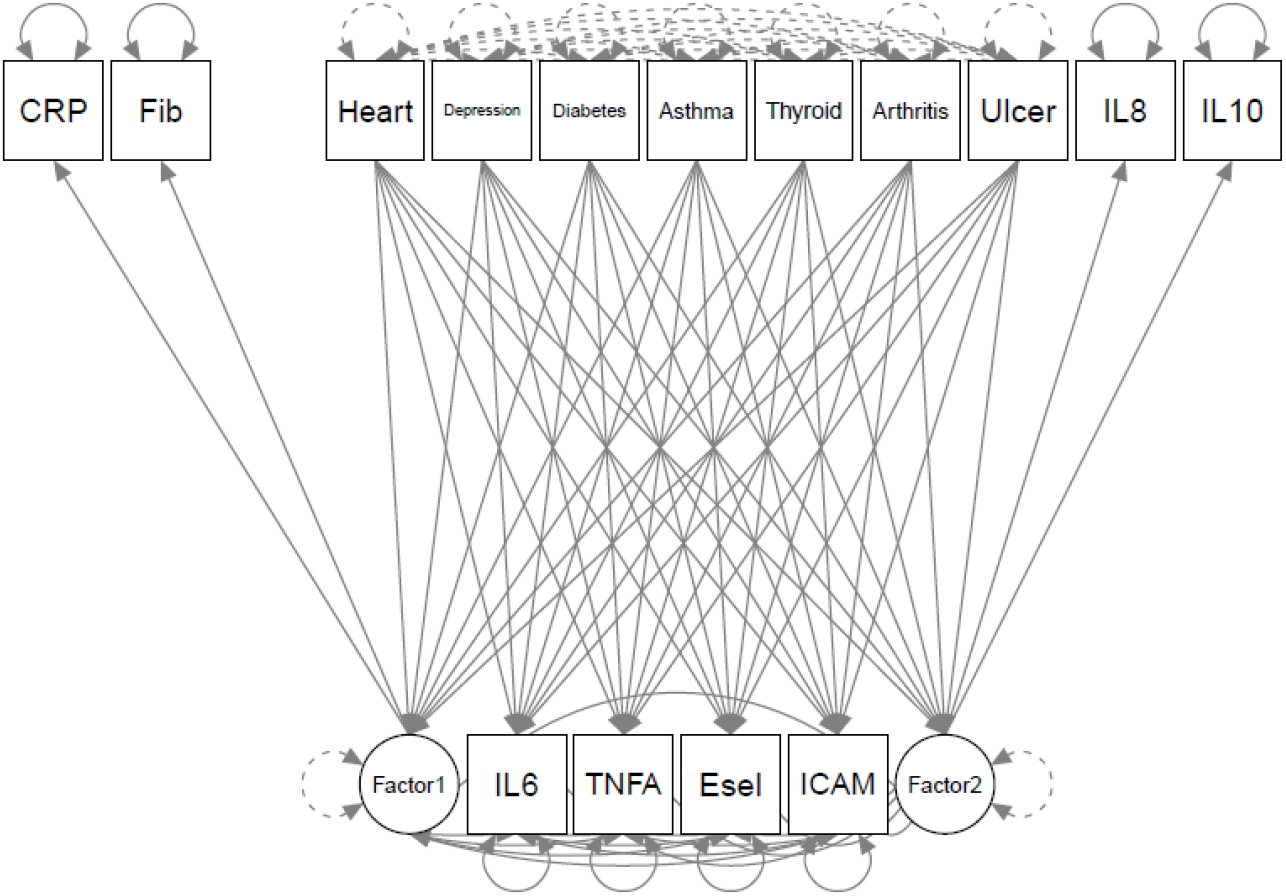
Health Predictors of Empirically-identified Structure.

**Figure 2b.**
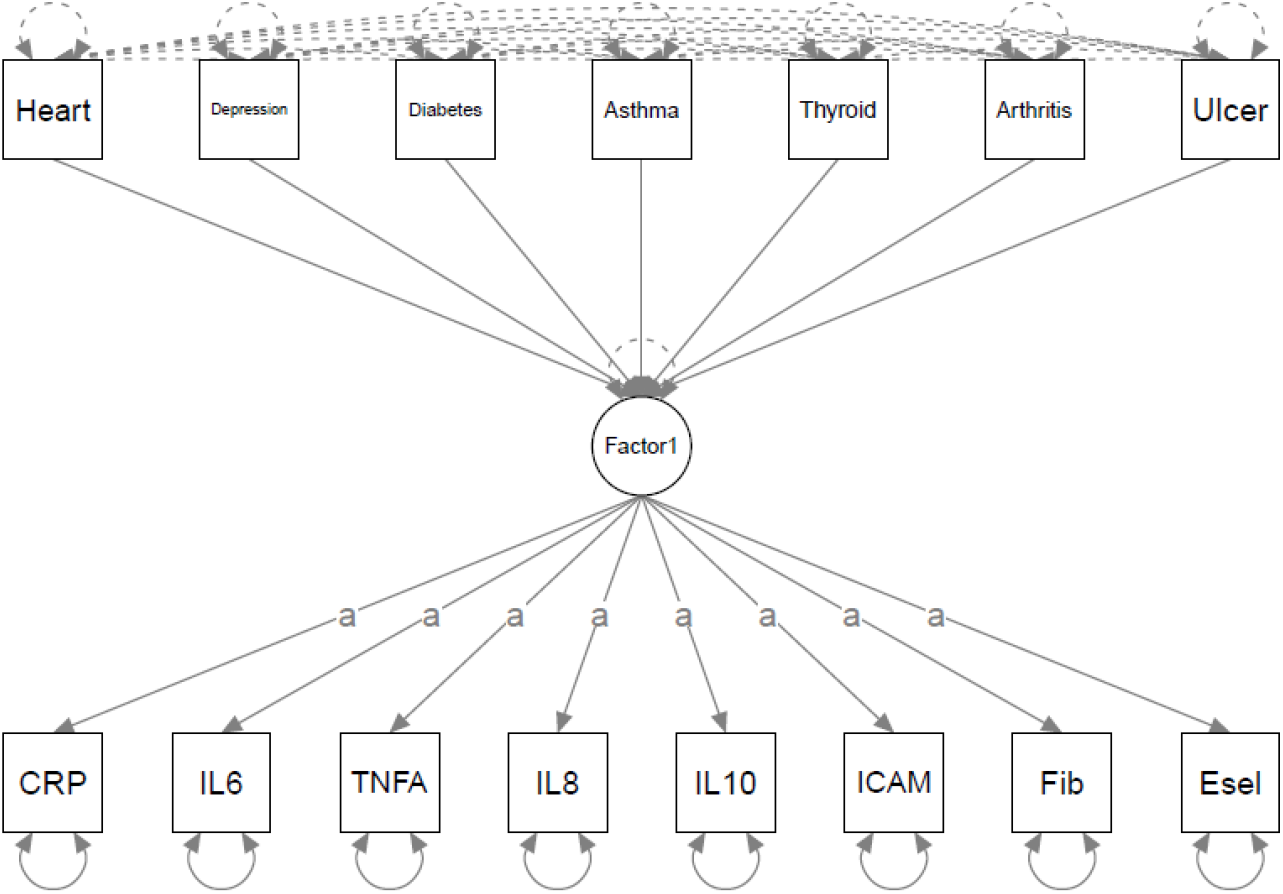
Health Predictors of A Priori/Tau-Equivalent Factor. Note: “a” indicates loadings constrained to equality. CRP = C-reactive Protein, Fib=fibrinogen, IL = interleukin, TNFA= Tumor Necrosis Factor-α, ICAM = Intracellular Adhesion Molecule-1, E-sel = E-selectin.

**Figure 2c.**
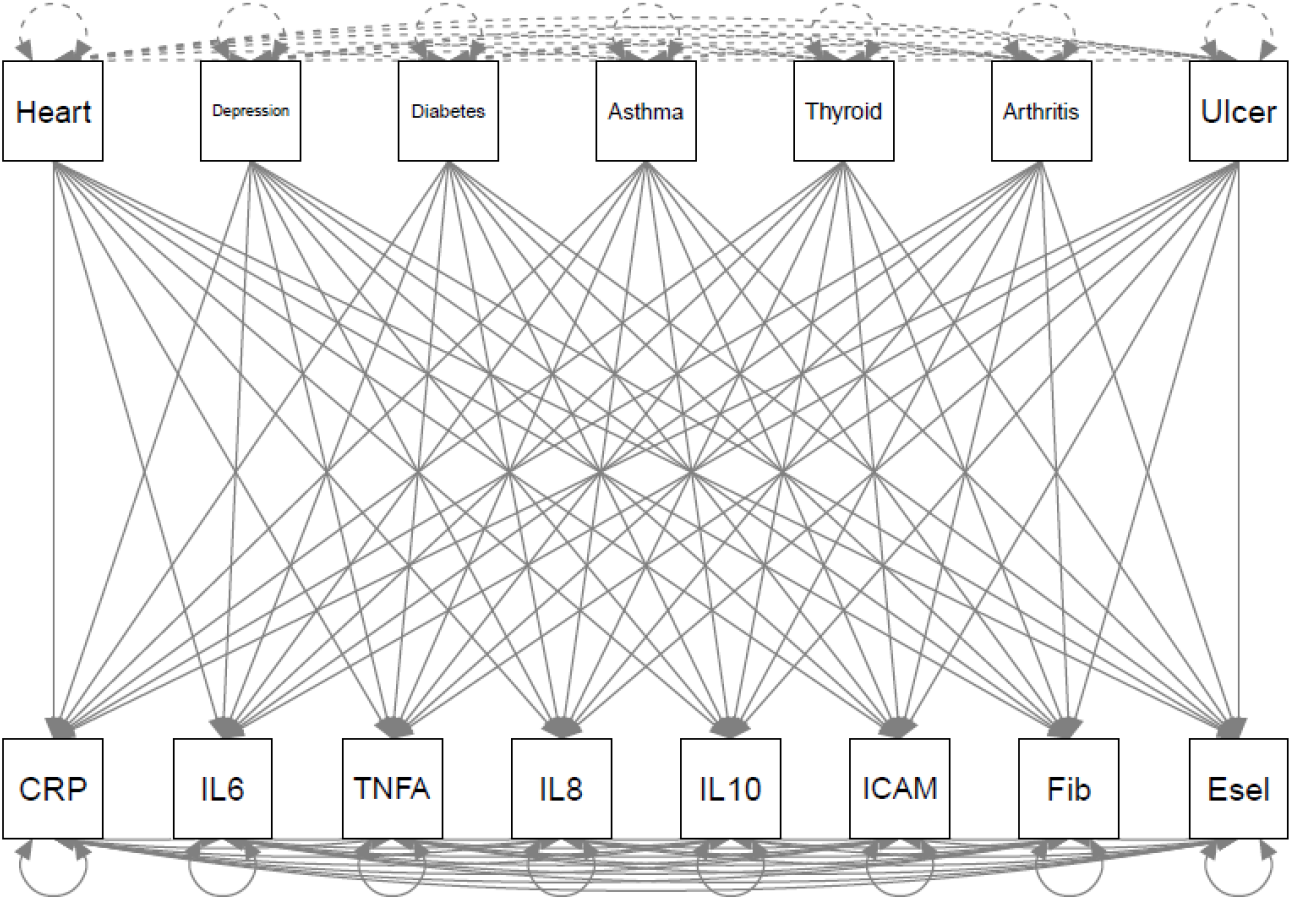
Health Predictors of Individual Proteins. Note: CRP = C-reactive Protein, Fib=fibrinogen, IL = interleukin, TNFA= Tumor Necrosis Factor-α, ICAM = Intracellular Adhesion Molecule-1, E-sel = E-selectin.

Because the individual protein model was just-identified, only AIC and BIC were able to be estimated. Fortunately, these statistics allow direct comparison to other models with the same variables, the primary aim of this study. The empirically-identified model (Table 3: AIC/BIC = 45511.005/45869.375) and individual protein models (AIC/BIC = 45493.632/45959.047) both outperformed the “a priori”/tau-equivalent model (AIC/BIC = 46447.441/46559.141). Compared to the individual protein model, the empirically-identified model had better BIC, but worse AIC, suggesting no clear answer as to which fit the data better.

**Table 3.** Fit Statistics of Different Inflammatory Models.

| | Robust<br>$\chi^2$ | Robust<br>$\chi^2$ df | Robust<br>$p_{\chi^2}$ | Robust<br>CFI | Robust<br>RMSEA | Robust<br>90% CI<br>RMSEA | SRMR | AIC | BIC |
| --- | --- | --- | --- | --- | --- | --- | --- | --- | --- |
| **MIDUS-R CFAs (N = 849)** |  |  |  |  |  |  |  |  |  |
| A priori/Tau-equivalent | 246.150 | 27 | <.001 | .089 | .184 | .163-.206 | .160 | 50864.267 | 50944.916 |
| Empirically-identified | 19.101 | 2 | <.001 | .966 | .068 | .042-.097 | .022 | 25598.958 | 25655.887 |
| **MIDUS-R SEMs with Medical Criteria (N = 776)** |  |  |  |  |  |  |  |  |  |
| A priori/Tau-equivalent | 1231.540 | 76 | <.001 | .077 | .133 | .126-.139 | .105 | 46447.441 | 46559.141 |
| Empirically-identified | 109.019 | 23 | <.001 | .956 | .053 | .043-.063 | .021 | 45511.005 | 45869.375 |
| Individual proteins | .000 | 0 | N/A | 1.000 | .000 | .000 | .000 | 45493.632 | 45959.047 |

Note: $\chi^2$ = Chi-squared, df = degrees of freedom, $p$ = $p$ -value, CFI = Comparative Fit Index, RMSEA = Root Mean Square Error of Approximation, CI = Confidence Interval, SRMS = Standardized Root Mean Square Residual, AIC = Akaike Information Criterion, BIC = Bayesian Information Criterion, MIDUS-R = Midlife in the United States-Refresher, CFA= Confirmatory Factor Analysis, SEM= Structural Equation Model, IL-6 = Interleukin-6

**Table 4.** Protein Loadings.

|  | MIDUS-2 EFA<br>(N = 1231) |  | MIDUS-R CFA<br>(N = 849) |  |
| --- | --- | --- | --- | --- |
|  | Factor 1 | Factor 2 | Factor 1 | Factor 2 |
| C-reactive Protein | .77 |  | **.84\***<br>**90% CI = .515-1.161** |  |
| Interleukin-6 |  |  |  |  |
| Tumor Necrosis Factor-α | | | | |
| Interleukin-8 |  | .76 |  | .48*<br>90% CI = .36-.60 |
| Interleukin-10 |  | .43 |  | **.51**<br>**90% CI = .09-.96** |
| Fibrinogen | .63 |  | **.51**<br>**90% CI = .33-.68** |  |
| Intercellular Adhesion Molecule-1 |  |  |  |  |
| E-selectin |  |  |  |  |

Note: \* = constrained to equality (note that these are standardized estimates so they will no longer be equal). Proteins not substantively loaded onto the factor were not depicted. CFA confidence intervals that include the original EFA estimate are **bolded**. MIDUS = Midlife in the United States, EFA = Exploratory Factor Analysis, CFA = Confirmatory Factor Analysis, CI = Confidence Interval

Because of unacceptable model fit (Table 3), the associations between the health conditions and the “a priori”/tau-equivalent inflammation variable are not reported. The associations between the medical conditions and a) the empirically-identified factors and b) the individual proteins are reported in Table 5. In the model with the empirically-identified factors and the individual proteins that did not load onto a factor, both empirically-identified inflammatory factors, IL-6, and E-selectin were significantly associated with a history of diabetes (Factor 1 β = .101, *b* =.769, *SE* = .187, *p* < .001; Factor 2 β = .164, *b* =.555, *SE* = .175, *p* = .002; IL-6 β = .066, *b* =.385, *SE* = .180, *p* = .032; E-selectin β = .179, *b* =10.791, *SE* = 2.626, *p* < .001). Additionally, Factor 1 was associated with depression (β = .101, *b* =.224, *SE* = .112, *p* = .046), Factor 2 was associated with arthritis (β = .134, *b* =.294, *SE* = .122, *p* = .016), and ICAM-1 was associated with heart disease (β = −.055, *b* =-29.959, *SE* = 11.492, *p* = .009); however, these three results (43% of significant results) were no longer significant after family-wise Benjamini-Hochberg false-discovery rate corrections (BH-FDR). Families were grouped by independent variable (i.e., health conditions). Thus, these relations between health conditions and inflammatory variables were corrected for six tests and the relations between health conditions and individual proteins below were corrected for eight.

**Table 5.**
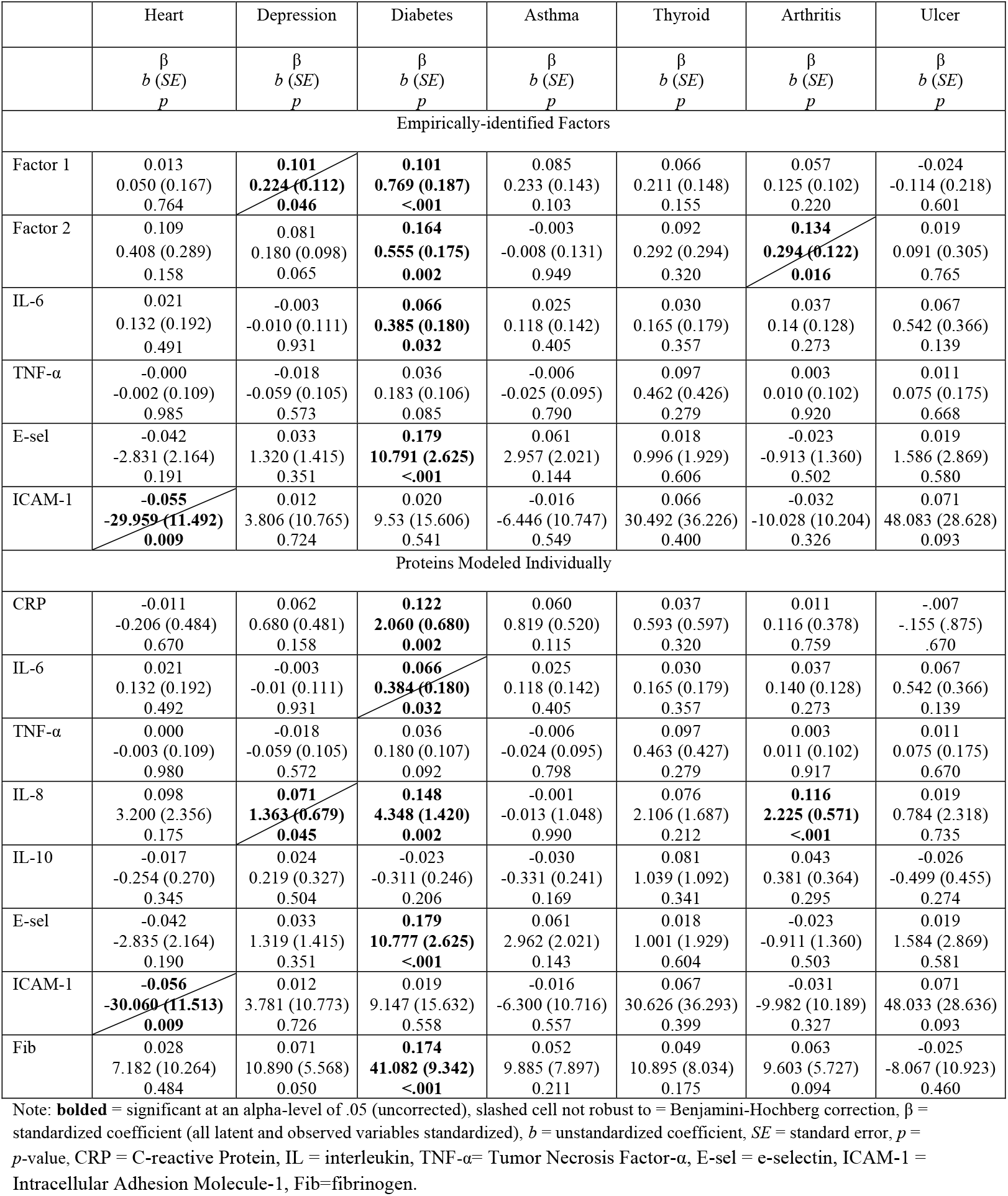
Health Conditions Predicting Inflammatory Outcomes (N = 776)

In the model with the medical conditions predicting individual proteins, diabetes was significantly associated with CRP (β = .122, *b* = 2.060, *SE* = .680, *p* = .002), IL-8 (β = .148, *b* =4.348, *SE* = 1.420, *p* = .002), E-selectin (β = .179, *b =10.777, SE* = 2.625, *p* < .001), and fibrinogen (β = .174, *b* = 41.082, *SE* = 9.342, *p* < .001). Arthritis was associated IL-8 (β = .116, *b* = 2.225, *SE* = .571, *p* < .001). These five results were robust to BH-FDR corrections. The following three associations (38% of significant results), were no longer significant after BH-FDR corrections: diabetes was associated with IL-6 (β = .066, *b* =.384, *SE* = .180, p = .032), heart disease was associated with ICAM-1 (β = −.056, *b* =30.060, *SE* = 11.513, *p* = .009)., and IL-8 was associated with depression (β = .071, *b* =1.363, *SE* = .679, p = .045).

## Discussion

Most studies investigating inflammation in relation to medical or psychiatric illnesses test individual proteins as predictors and/or outcomes. This invites problems with multiple comparisons and creates a disconnect between theories about generalized inflammation and the analyses conducted. Alternatively, some studies (e.g., Chat et al., 2021; Moriarity et al., 2020; Tait et al., 2019; Vinhaes et al., 2021) use composite variables created without first investigating the appropriateness of this decision in what we describe as “a priori”/tau-equivalent composites. The present study sought to apply standard aggregate-building procedures to a set of eight inflammatory proteins to critically evaluate the statistical viability of this approach and compare it to empirically-identified inflammatory factors and individual proteins. The current analyses indicated that, out of the eight proteins available in the MIDUS datasets, two factors emerged. Specifically, CRP and fibrinogen loaded onto the first factor (Factor 1, interpreted to reflect the acute phase reaction) and IL-8 and IL-10 loaded onto a second factor (Factor 2, interpreted to reflect inflammatory processes (both pro- and anti-inflammatory) particularly associated with neutrophil activity). Across all models estimated, the “a priori”/tau-equivalent model did not fit the data well enough to be considered a valid approach. In fact, the empirically-identified structure and modeling individual proteins out-performed this approach in all comparisons. Further, reliability estimates for all latent variables (both “a priori”/tau-equivalent and “empirically-identified”) were poor.

Fit criteria diverged on whether the empirically-identified structure fit the model with medical variables better than modeling the proteins individually. Specifically, BIC supported the empirically-identified structures and AIC supported individual markers. Unfortunately, because all variables in the individual protein model were observed variables (resulting in a “just-identified” model), it is impossible to evaluate other fit statistics. Both sets of analyses resulted in primarily the same conclusion (inflammation was most robustly associated with a history of diabetes). However, it is worth note that a larger proportion of results were robust to BH-FDR corrections (despite a larger number of total analyses) in the individual protein model compared to the empirically-identified factors model. Thus, there might be a level of specificity in associations that were washed out when using the latent variable models due to decreased signal-to-noise ratios, nullifying any potential benefit of combining proteins into composites with respect to multiple comparisons. However, these findings are far from conclusive and future research is necessary. Additionally, careful consideration should be given to whether variable aggregation is a theoretically-appropriate course of action (e.g., whether it is theoretically sound to aggregate often pleotropic proteins into composites or whether specific biomarkers are theorized to drive effects rather than generalized inflammation). Further, this study adopted a latent modeling approach to form factors; it is possible that other types of modeling (e.g., causal indicator models, network models, multiple-indicator multiple-cause (MIMIC) models) may be more suitable for modeling inflammatory processes.

Interestingly, the two proteins that loaded onto Factor 1 (CRP and fibrinogen) also loaded onto the same factor in prior research in a community sample of older adults (Egnot et al., 2018), a sample of adults with unstable angina pectoris (Koukkunen et al., 2001), a samples of patients with insulin resistance syndrome (Sakkinen et al., 2000), and a sample of patients with acute coronary syndrome (Tziakas et al., 2007). Also consistent with this study, ICAM-1 was measured, but did not load onto this factor in Egnot et al. (2018) or Tziakas et al. (2007). Similarly, TNF-α also was measured, but did not load onto this factor in Koukkunen et al. (2001). Consequently, there is evidence that this grouping is not an artifact of the specific array of proteins available in this study or other study-specific methods, demonstrating both internal replicability in this study and external replicability with previous studies. It is worth note that IL-6 loaded onto this factor in several studies (Egnot et al., 2018; Koukkunen et al., 2001) and a previous version of this manuscript. Notably, both papers and the previous version of this manuscript used factor analyses in which normality was handled by transforming values (or didn’t report on how non-normality was handled at all) instead of running models robust to non-normality, which might drive this difference and highlight the importance of using robust estimation techniques with skewed data. Future research is needed to test whether the structure of inflammatory proteins is stable in more dynamic contexts (e.g., across acute or chronic stressors) and over time (e.g., weeks or months).

It is critical to underscore the pleotropic nature of many inflammatory proteins and the complexity of inflammatory processes. For example, IL-6, commonly conceptualized as a “pro-inflammatory” protein, has several anti-inflammatory functions that are most likely dependent on classic-signaling via the membrane-bound non-signaling α-receptor IL-6R (Scheller et al., 2011). Given the contextual functioning of the immune system, shared variance analyses might be more informative in the context of acute inflammatory activity compared to the resting data used in this study. Truly, due to the complexity of this system, comparable model fit to the empirically-indicated factor structure, poor factor reliability, and greater robustness of results to multiplicity correction, it might be more appropriate to analyze specific proteins and plan around the inherent limitations of doing so. However, should the decision be made to use aggregate variables of inflammation, the results of this study consistently discourage the use of aggregates in which all proteins are equally weighted (referred to as an “a priori”/tau-equivalent composite throughout). Instead, standard aggregate measure-building procedures should be utilized to inform data aggregation (for more detailed concerns about the use of straight sum scoring to create aggregates when not all component variables are equally associated with the construct of interest, see McNeish & Wolf (2020)). Until a more comprehensive understanding of how to model inflammation is achieved, researchers may benefit from simultaneously testing multiple scales of inflammatory measurement (Moriarity & Alloy, 2020) or focusing on individual proteins.

This study has several important strengths. First, two sizeable datasets were used allowing for replication. Second, the replication sample was specifically designed for replication analyses, maximizing comparability of methods. Third, the fact that both samples were community samples maximizes generalizability to other non-medical samples and expands upon the scant work on the dimensionality of inflammatory proteins that has primarily used medical samples. Still, ideal modeling techniques could differ for certain medical populations and additional work is needed to thoroughly vet the generalizability of the structure of inflammatory proteins. Additionally, this study used methods robust to non-normal data structures, whereas many studies in this field transform the data to fit the models (resulting in raw data that is not truly representative of the sample under study).

However, this study must be interpreted in the context of several limitations. First, although eight proteins are more than many studies, there are a lot of inflammatory markers that were not included (e.g., IL-1β, T-cells, B-cells). Thus, the empirically-identified structure might look different when a broader array/different proteins are used. However, the finding that CRP and fibrinogen loaded onto one factor (which did not include ICAM-1 or TNF-α) is consistent with previous investigations of the dimensionality of inflammation in medical and community samples (e.g., Egnot et al. (2017)), supporting the generalizability of this factor. Second, the number of identifiable latent factors is constrained by the number of indicator variables. Consequently, a study with more than eight indicator variables might find a more multidimensional factor structure. Relatedly, factors with less than three indicators (as was the case for both observed factors in this study) are generally not adequately reliable and either involve constraining parameters to equality to identify a model solution (as was done in the confirmatory factor analyses) or can be locally unidentified despite global model identification (as is the case in the models with health criterion). Third, the ability to test these factors in the context of acute inflammatory activity (e.g., endotoxin exposure, acute stress task) would provide great insight into their biological plausibility. Additionally, although several studies using community and clinical samples have found CRP and fibrinogen to load onto the same factor, it would be insightful for future studies to directly test the comparability of inflammatory structure as a function of medical status using measurement invariance testing. Finally, the medical criterion variables were measured via self-report of a *history* of a diagnosis. These results would be more informative if active medical conditions were known.

## Conclusion

This study used standard aggregate measure building procedures to investigate the structure of eight inflammatory proteins. Results firmly indicate that an “a priori”/tau-equivalent aggregate in which all inflammatory proteins equally load onto a single dimension did not reflect the data accurately enough to warrant use in research. Both a two-factor empirically-identified structure and individual proteins were preferrable. Although there is no conclusive evidence about which of these two options are preferrable, given comparable model fit, similar conclusions in tests of predictive validity, results that were more robust to correcting for multiple comparisons, unacceptable latent factor reliability, and ease of interpretation, we recommend researchers either analyze individual proteins or explore multiple levels of inflammatory measurement (e.g., factors and individual proteins) until more work is done to explore the empirical and theoretical support for inflammatory factors. For example, direct comparison of inflammatory structure in samples with different medical statuses and tests of structural invariance across acute inflammatory reactions would be important studies to extend this line of inquiry. To facilitate other researchers in designing studies and analyzing data, indices of model fit and factor reliability were described as appropriate. By building a strong physiometric foundation, research on inflammation can become more replicable, cost-effective, and clinically-impactful (Moriarity, 2021).

## Acknowledgments

Daniel P. Moriarity was supported by National Research Service Award F31 MH122116 and an APF Visionary Grant. Lauren B. Alloy was supported by National Institute of Mental Health R01 MH101168. Lauren M. Ellman was supported by the National Institute of Mental Health R01 MH096478 and R01 MH11854. Thomas M. Olino, was supported by the National Institute of Mental Health R01 MH107495. The MIDUS project was supported by an award from the National Institute on Aging (U19 AG051426) and sample collection at the 3 Clinical and Translational Research Units enabled by support from M01-RR023942, M01-RR00865, and U01TR000427.

---

**Supplemental Table 1.** Bivariate Spearman Correlations of Proteins in MIDUS-2

| Protein | 1. | 2. | 3. | 4. | 5. | 6. | 7. | 8. |
| --- | --- | --- | --- | --- | --- | --- | --- | --- |
| 1. CRP | — |  |  |  |  |  |  |  |
| 2. IL-6 | .52 | — |  |  |  |  |  |  |
| 3. TNF-α | .23 | .34 | — | | | | | |
| 4. IL-8 | .02 | .12 | .21 | — |  |  |  |  |
| 5. IL-10 | .08 | .15 | .35 | .20 | — |  |  |  |
| 6. Fibrinogen | .53 | .41 | .11 | .05 | .04 | — |  |  |
| 7. E-selectin | .15 | .16 | .09 | .09 | .10 | .11 | — |  |
| 8. ICAM-1 | .19 | .18 | .29 | .07 | .13 | .12 | .05 | — |

Note: N = 1231. CRP = C-reactive protein; IL- = Interleukin; TNF-α = tumor necrosis factor alpha; ICAM= intracellular adhesion molecule

**Supplemental Table 2.** Bivariate Spearman Correlations of Proteins in MIDUS-R

| Protein | 1. | 2. | 3. | 4. | 5. | 6. | 7. | 8. |
| --- | --- | --- | --- | --- | --- | --- | --- | --- |
| 1. CRP | — |  |  |  |  |  |  |  |
| 2. IL-6 | .57 | — |  |  |  |  |  |  |
| 3. TNF-α | .23 | .37 | — | | | | | |
| 4. IL-8 | .06 | .17 | .24 | — |  |  |  |  |
| 5. IL-10 | .04 | .14 | .39 | .13 | — |  |  |  |
| 6. Fibrinogen | .57 | .45 | .18 | .13 | .01 | — |  |  |
| 7. E-selectin | .24 | .28 | .29 | .06 | .17 | .18 | — |  |
| 8. ICAM-1 | .16 | .15 | .24 | .03 | .18 | .13 | .37 | — |

Note: N = 849. CRP = C-reactive protein; IL- = Interleukin; TNF-α = tumor necrosis factor alpha; ICAM= intracellular adhesion molecule

